# Peptide fusion improves prime editing efficiency

**DOI:** 10.1101/2021.09.22.461415

**Authors:** Minja Velimirovic, Larissa C. Zanetti, Max W. Shen, James D. Fife, Lin Lin, Minsun Cha, Ersin Akinci, Danielle Barnum, Tian Yu, Richard I. Sherwood

## Abstract

Prime editing enables search-and-replace genome editing but is limited by low editing efficiency. We present a high-throughput approach, PepSEq, to measure how fusion of 12,000 85-amino acid peptides derived from human DNA repair-related proteins influences prime editing efficiency. We show that peptide fusion can enhance prime editing, prime-enhancing peptides combine productively, and a top dual peptide-prime editor increases prime editing significantly in multiple cell lines across dozens of target sites.

---

Prime editing is a CRISPR-based *‘search-and-replace’* technology that mediates targeted insertions, deletions, and all possible base-to-base conversions in mammalian cells in the absence of double-stranded breaks (DSBs) or donor DNA templates^1^. The prime editing enzyme (PE2) consists of SpCas9-nickase fused to an engineered reverse transcriptase (RT). PE2 is recruited to a target site by a prime editing guide RNA (pegRNA) which, in addition to a standard genome-targeting spacer and SpCas9-binding hairpin, contains a 3’ sequence that acts as a template for the fused RT to synthesize a programmed DNA sequence on one of the nicked DNA strands. When cellular DNA repair machinery repairs the broken strand, this RT-extended flap competes with the unedited flap, and the edited sequence sometimes replaces the original sequence in the genome^1,2^.

Because of its versatility, prime editing has enormous potential in improving understanding of the effects of genetic changes on cellular and organismal function. However, prime editing is limited by low efficiency. While editing efficiency is dependent on the experimental system, a survey of lentiviral PE2 efficiency at thousands of sites found that PE2 rarely leads to installed edits in >20% of alleles^3^. Analysis of features associated with prime editing efficiency at these thousands of loci found that the strongest correlate is DeepSpCas9 score^3,4^, suggesting that prime editing is limited by the interaction strength between the SpCas9-pegRNA complex and the target locus. Optimization of pegRNA features^3^, induction of a distal nick on the opposite strand (designated PE3)^1^, and pairing overlapping pegRNAs^5^ have been found to improve prime editing efficiency, yet low efficiency remains an issue in deployment of prime editing.

Here, we screened a library of 12,000 85-amino acid peptides derived from DNA repair proteins to identify peptides that improve prime editing efficiency when appended to the N-terminus of PE2. Peptide and protein fusion is a well-established method of modulating genome editing outcomes^6–8^. While scalable, sensitive protein fusion screening remains challenging, high-throughput oligonucleotide library synthesis enables screening of highly diverse peptide fusion constructs. Reasoning that peptides derived from DNA repair-related proteins may encode domains capable of altering prime editing efficiency, we designed a library of 85-amino acid peptides comprising complete 2X tiling of 417 DNA repair-related proteins^9,10^ and 29 housekeeping genes as controls (Supplementary Table 1). We also included 5,458 DNA repair-related mutant peptides with all possible S-->E and T-->E phosphomimetic substitutions. This library of 12,000 oligos was cloned N-terminal to a 33-amino acid XTEN linker followed by PE2 in a vector allowing Tol2 transposon-mediated genomic integration (Fig. 1a)^11^.

**Figure 1.**
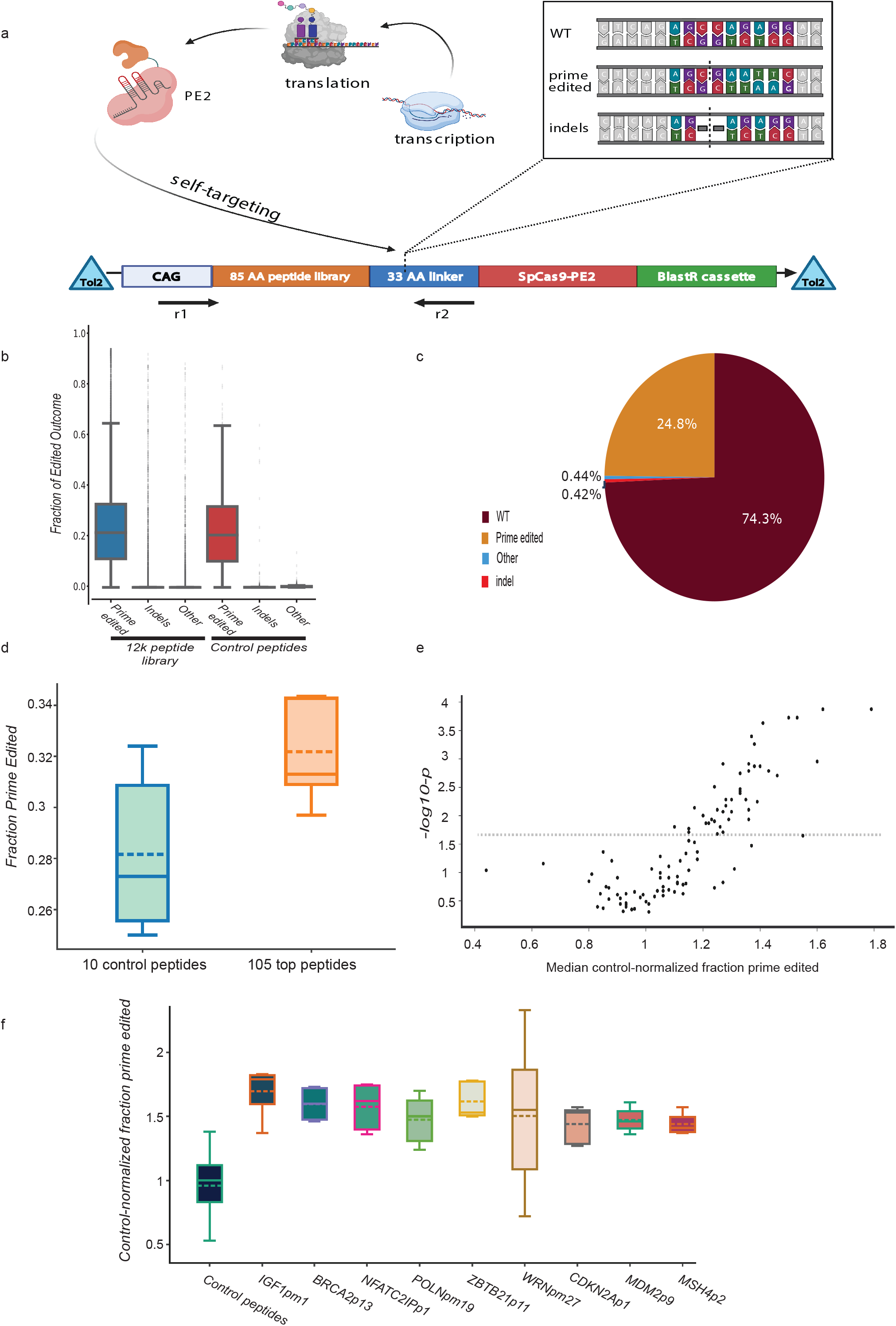
The high-throughput Peptide Self-Editing sequencing assay (PepSEq) identifies peptides capable of increasing prime editing efficiency. (a) In PepSEq, a library of peptides from human DNA repair-related genes is cloned N-terminal to SpCas9-PE2 separated by a linker and integrated into cells at one copy per cell. Cells are subsequently treated with a pegRNA targeting the linker sequence that installs a fixed edit. Paired-end genomic DNA NGS of the peptide sequence and the editing site allows calculation of prime editing outcomes in high throughput. (b) Observed prime editing outcome frequencies for a 12,000-peptide PepSEq screen. Box plot indicates median and interquartile range, and whiskers indicate extrema. (c) Overall distribution of prime editing outcome frequencies across all peptides and replicates. (d) Comparison of prime edited allele fraction for 105 DNA repair-related peptides vs. 10 housekeeping control peptides from a 115-peptide PepSEq screen. (e) Volcano plot showing control-normalized prime editing fold change (x-axis) vs. vs. –log_10_ p-value (y-axis) from 115-peptide PepSEq screen. (f) Comparison of control-normalized prime edited allele fraction for nine top peptides and all control peptides from a 115-peptide PEPSeq screen.

To enable quantitative evaluation of peptide-PE2 editing efficiency in high-throughput, we devised the Peptide Self-Editing sequencing assay (PepSEq). We designed a self-targeting pegRNA that introduces a 6-nt mutation (CCTCTG-->GAATTC) in the peptide-adjacent linker sequence (sgPE-linker). Following Tol2-mediated genomic integration of a single peptide-PE2 library member per cell ^12^, cells are treated in pooled format with sgPE-linker. To evaluate prime editing efficiency in pooled format, we perform paired-end nextgen sequencing (NGS), mapping peptide-PE2 identity and genotypic outcome for each self-targeted allele (Fig. 1a). We performed initial PepSEq screens in mouse embryonic stem cells (mESCs) because they are a non-transformed cell line not known to possess DNA repair defects, in contrast to other common models such as HEK293T, which is known to lack mismatch repair capacity^13^.

We performed three biological replicates of peptide-PE2 integration, each followed by two biological replicates of sgPE-linker addition, collecting >30M NGS reads for each replicate and 10M reads prior to sgPE-linker treatment. We developed a computational pipeline to filter and analyze the NGS data (Online Methods), removing peptide-PE2s with <100 total reads in a given replicate from analysis. In total, 16-28% of all alleles were prime edited, 0.3-0.7% were indels, and nearly all remaining reads were unedited (Fig. 1b-c, Supplementary Fig. 1, Supplementary Table 2). Due to the low frequency of indels and other alleles, we focused analysis on prime editing efficiency for each peptide-PE2. Replicates that shared peptide-PE2 integration had strong consistency in prime editing efficiency (R = 0.42-0.66) while those with distinct peptide-PE2 integration had negligible consistency (R = 0.01-0.06) (Supplementary Fig. 1,). It is not surprising to obtain poor replicate consistency in a high-throughput screen in which the majority of library members are expected to be inert, so we analyzed the three combined replicates with independent peptide-PE2 integration using a beta-binomial model to identify 105 top candidate prime editing-enhancing peptide-PE2s (Online Methods).

To obtain higher-resolution data on this set of candidate peptide-PE2s, we cloned a sub-library with these 105 peptide-PE2s and 10 control peptide-PE2s, performing PepSEq in five biological replicate peptide-PE2 integrations in mESCs each with >1M NGS reads. Replicate consistency was much higher (R = 0.25-0.52, Supplementary Fig. 2, Supplementary Table 3), and the 105 candidate peptide-PE2s as a group gave 15% higher prime editing than control peptide-PE2s (P<0.0001, Fig. 1d), indicating that peptide fusion can improve prime editing efficiency. This screen identified 44 peptide-PE2s that significantly improve prime editing efficiency (FDR = 0.05), increasing prime editing efficiency up to 70% (Fig. 1e-f). The proteins from which these peptides are derived are not robustly enriched in any particular DNA repair pathway, and none encompass known functional domains that appear related to hypothesized prime editing mechanisms^1^. This result additionally indicates that the 12,000-peptide PepSEq screen was able to flag true hits in spite of noise, a finding supported by the fact that the 44 peptide-PE2s that significantly increase prime editing in the smaller screen have appreciable replicate consistency in the 12,000-peptide screens (R = 0.17-0.39, Supplementary Fig. 2,).

To gain insight into how these peptides function, we next asked whether peptides that increase prime editing combine productively. We constructed a dual peptide-PE2 library in which nine top candidate peptides and one control peptide were combined in all 100 possible combinations separated by an eight amino acid linker (Fig. 2a), and we performed 10 biological replicate PepSEq screens in mESCs and two replicates in HCT-116 colorectal carcinoma cells. These replicates were highly concordant within and between cell lines (mESC median R = 0.62, HCT-116 R = 0.47, mESC vs. HCT-116 median R = 0.32, Supplementary Fig. 3, Supplementary Table 4), and 79 of the 81 candidate dual peptide-PE2s gave significantly higher prime editing efficiency than the control-control pair in mESCs (Fig. 2b, Supplementary Fig. 3).

**Figure 2.**
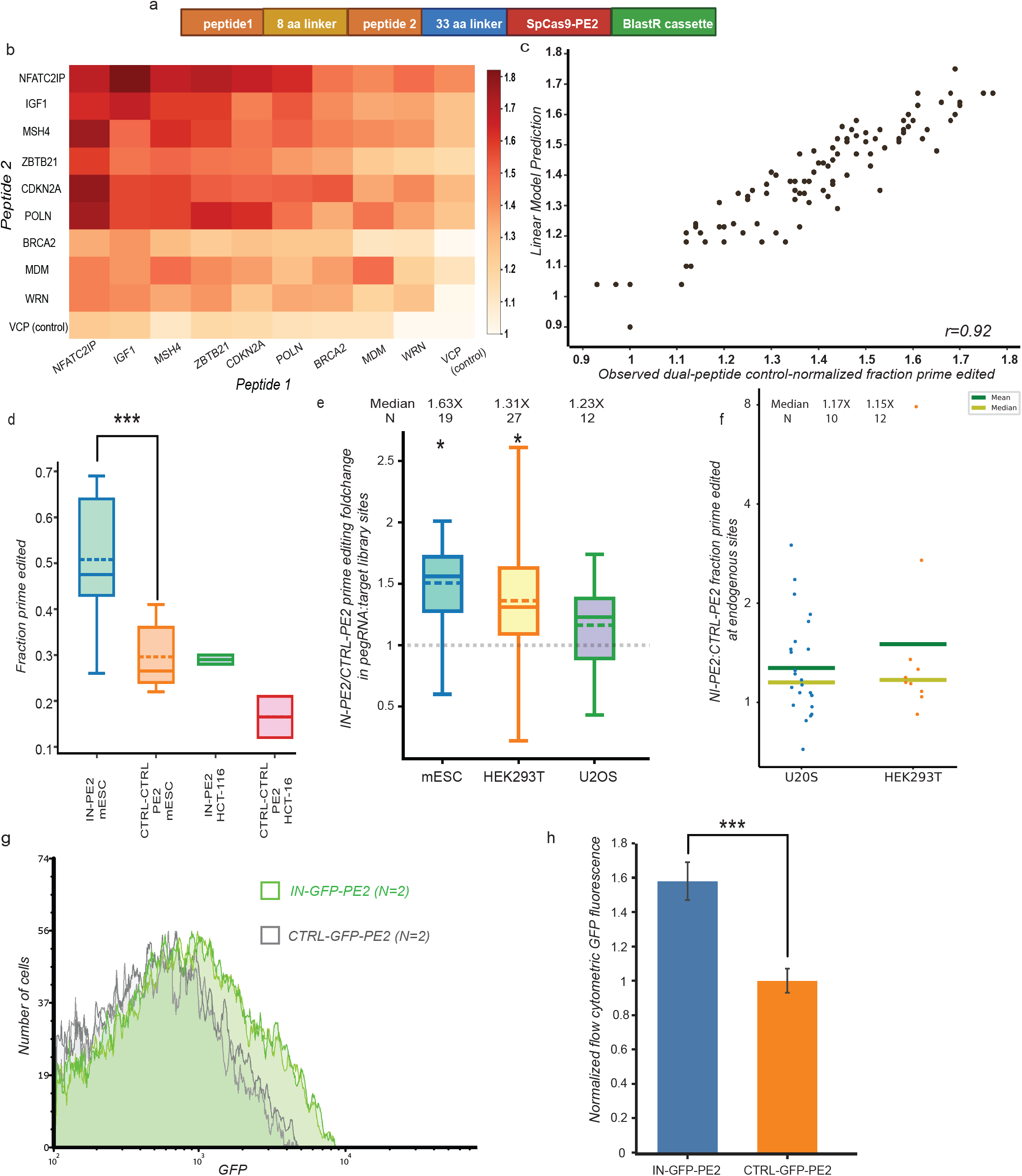
A dual-peptide-PE2 displays improved prime editing efficiency across dozens of loci in four cell lines. (a) Construct design for a dual-peptide PepSEq library with all pairs of 10 peptides. (b) Comparison of control-normalized prime edited fraction for 81 dual-peptide pairs. (c) Comparison of control-normalized dual-peptide prime edited fraction predicted by a linear model vs. observed median prime edited fraction. (d) Comparison of prime edited fraction for IN-PE2 vs. control-control-PE2 in mESC and HCT-116 dual-peptide screen replicates. (e) Comparison of median control-normalized prime edited fraction for IN-PE2 in the 100-target library across three cell lines. (f) Comparison of prime edited fraction for IN-PE2 vs. CTRL-PE2 at a set of six endogenous sites in HEK293T and U2OS. (g) Flow cytometric GFP fluorescence intensity for two representative replicates of IN-GFP-PE2 vs. CTRL-GFP-PE2 in mESCs. (h) Comparison of IN-GFP-PE2 vs. CTRL-GFP-PE2 control-normalized flow cytometric GFP fluorescence levels. N = 5.

To explore relationships between peptides, we asked whether dual-peptide-PE2 activation could be predicted by a linear model assuming consistent peptide-specific influences on prime editing. We find high consistency among observed prime editing and expected prime editing under additive assumptions by linear estimates (r = 0.92) (Fig. 2c). The high accuracy of the linear model suggests that each peptide has independent (not redundant or synergistic) effects on prime editing, either through interacting with distinct pathways or providing a fixed advantage in protein stability or DNA binding.

Eight of the nine dual peptide-PE2s with highest prime editing included an N-terminal peptide from NFATC2IP (NFATC2IPp1), and the dual-peptide with highest median prime editing in mESC and HCT-116 (median 1.77X control-control in mESC, 1.88X in HCT-116) pairs NFATC2IPp1 with a phosphomimetic peptide from IGF1 (IGF1pm1) (Fig. 2d). These two peptides induce the strongest increases in prime editing in the single-peptide-PE2 screen (Fig. 1c) and in the linear model of dual-peptide-PE2 screen (Supplementary Table 4), providing rationale to pursue IGFpm1-NFATC2IPp1-PE2 (IN-PE2) as an improved prime editor.

To ask whether IN-PE2 increases prime editing efficiency across a larger collection of target sites, we designed a lentiviral library comprising 100 pegRNA-target pairs spanning a range of edit types and predicted editing efficiencies^3^ (Methods). After stable integration of this library in three human and mouse cell lines (mESC, HEK293T, U2OS), we treated cells with either IN-PE2 or a control PE2 containing the 5’ linker sequence but lacking an N-terminal peptide (CTRL-PE2). Among the targets with sufficient library representation and editing, IN-PE2 yielded significantly higher prime editing than CTRL-PE2 in all three cell lines (median 1.63X in mESC at 19 sites, 1.31X in HEK293T at 27 sites, 1.23X in U2OS at 12 sites) Fig. 2e, Supplementary Fig. 4, Supplementary Table 5). The results indicate that IN-PE2 leads to a consistent increase in prime editing efficiency across a variety of targets (Supplementary Fig. 4). We performed pegRNA-target library experiments with six additional peptide-PE2s with one to three top candidate peptides fused to PE2, finding IN-PE2 to display the most robust prime editing of these seven peptide-PE2s (Supplementary Fig. 4). The consistent increase in prime editing efficiency in cell lines without known DNA repair defects (mESC, U2OS) and those with known deficiencies in mismatch repair (HEK293T^13^, HCT-116^14^) suggests that IN-PE2 is unlikely to function through interaction with mismatch repair machinery.

We next asked whether IN-PE2 increases prime editing at endogenous genomic loci. We transduced HEK293T and U2OS with a pool of six lentiviral pegRNAs encoding missense variants in exon 8 of the NF2 gene, each in two biological replicates. We subsequently treated cells with CTRL-PE2 or IN-PE2 and performed genomic DNA NGS to determine editing outcomes. IN-PE2 treatment led to significantly increased prime editing efficiency in both cell lines across the six loci (median 1.17X in HEK293T, 1.15X in U2OS, p<0.01 in each cell line) (Fig. 2f, Supplementary Fig. 5). There was variability in the magnitude of increased prime editing across sites, but all six loci had increased prime editing in each cell line, suggesting that IN-PE2 consistently increases prime editing efficiency at native genomic targets.

To gain insight into the mechanism by which IN-PE2 increases prime editing efficiency, we constructed IN-GFP-PE2 and CTRL-GFP-PE2 fusions to address whether IN-PE2 increases the amount of protein in cells. We found that cells possess 1.58X the amount of IN-GFP-PE2 as CTRL-GFP-PE2 (N=5, p<0.0001, Fig. 2g, Supplementary Fig. 6). Cycloheximide timecourse experiments show that IN-GFP-PE2 and CTRL-GFP-PE2 are degraded at a similar rate (Supplementary Fig. 6), altogether suggesting that the NI peptides increase either transcription or translation of the PE2 enzyme and offering a plausible explanation for the increased activity of IN-PE2.

In summary, through screening 12,000 peptide-PE2 fusion proteins using PepSEq, a sensitive, NGS-based self-editing platform, we identify a prime editor that consistently increases editing efficiency across dozens of targets in four human and mouse cell lines. Because prime editing applications are currently limited by editing efficiency, we anticipate that IN-PE2 will be a valuable tool in elucidating how DNA sequence influences genome function.

## Acknowledgments

The authors thank Mandana Arbab for technical assistance. The authors acknowledge funding from 1R01HG008754 (R.I.S.), 1R21HG010391 (R.I.S., C.A.C.), American Cancer Society (R.I.S.), American Heart Association (R.I.S.), Qatar Biomedical Research Institute (R.I.S.), and the São Paulo Research Foundation-FAPESP n° 2019/13813-6 and 2017/25009-1 (L.C.Z.). Figures created with BioRender.

## Author contributions

Conceptualization, Methodology, Writing – Original Draft and Writing – Reviewing and Editing: R.I.S., M.V., L.L., M.W.S., J.D.F. Investigation and Validation: R.I.S., M.V., L.C.Z., M.C., E.A., D.B., M.W.S., J.D.F., T.Y.; Software, Formal Analysis and Visualization: R.I.S., M.W.S., J.D.F., T.Y., M.V.; Funding Acquisition and Supervision: R.I.S.

## Competing interests

The authors report no competing interests.

## Online Methods

### Peptide library design

We designed 85-amino acid peptides covering all annotated human DNA repair proteins^16,17^, tiling by starting each peptide 45-amino acids after the prior peptide using a codon-optimized library design^15^. We also included mutant peptides with all possible S-->E and T-->E phosphomimetic substitutions. 147 wild-type peptides targeting 29 housekeeping genes were also included as controls. Unique 9-nt sequences were inserted in phosphomimetic peptides to facilitate sequence mapping for downstream analysis. The sequence design was performed with “seqinr” and “Biostrings” packages in R.

### Cell Culture

All cell lines were obtained from ATCC and were cultured in: McCoy’s 5A media (Thermo Fisher) + 10% FBS (Thermo Fisher) (U2OS, HCT-116); DMEM (Thermo Fisher) + 10% FBS (HEK293); mESCs were maintained on gelatin-coated plates feeder-free in mESC media composed of Knockout DMEM (Life Technologies) supplemented with 15% defined foetal bovine serum (FBS) (HyClone), 0.1 mM nonessential amino acids (NEAA) (Life Technologies), Glutamax (GM) (Life Technologies), 0.55 mM 2-mercaptoethanol (b-ME) (Sigma), 1X ESGRO LIF (Millipore), 5 nM GSK-3 inhibitor XV and 500 nM UO126. Cells were regularly tested for mycoplasma.

### Peptide library cloning and screening

The SpCas9-PE2-encoding sequence from pCMV-PE2^1^ (Addgene Plasmid #132775) was subcloned into the p2T-CAG-SpCas9-BlastR plasmid^18^ (Addgene Plasmid #107190) to create p2T-CAG-SpCas9PE2-5pLinker-BlastR (Addgene Plasmid #173066)

Specified oligonucleotide libraries were synthesized by Twist Bioscience (12,000-peptide) or IDT (115-peptide and dual-peptide) and were cloned into the NheI site of p2T-CAG-SpCas9PE2-5pLinker-BlastR through amplification with Q5® High-Fidelity DNA Polymerase (New England Biolabs) using primers Cas9NTLib_GA_fw and Cas9NTLib_GA_rv (see Supplementary Table) followed by ligation using the NEBuilder HiFi DNA Assembly Kit (NEB) for 1 h at 50 °C. Assembled plasmids were purified by isopropanol precipitation with GlycoBlue Coprecipitant (Thermo Fisher) and reconstituted in TE and transformed into NEB® 10-beta Electrocompetent *E. coli* (NEB). Following recovery, the library was grown in liquid culture in LB medium overnight at 37 °C and isolated by ZymoPURE™ II Plasmid Maxiprep Kit (Zymo Research). Library integrity was verified by restriction digest with AgeI (New England Biolabs) for 1 h at 37 °C, and library diversity was validated by Sanger sequencing sampling.

Mouse ESC cells were plated at ∼20-25% confluence onto 25-cm plates the day before transfection so that they reach ∼50-75% confluency on the day of transfection. For stable Tol2 transposon plasmid integration, cells were transfected using Lipofectamine 3000 (Thermo Fisher) following standard protocols, and equimolar amounts of Tol2 transposase plasmid and transposon-containing plasmid. To generate lines with stable Tol2-mediated genomic integration, selection with the appropriate selection agent at an empirically defined concentration (blasticidin, hygromycin, or puromycin) was performed starting 24 h after transfection and continuing for >1 week. In cases where sequential plasmid integration was performed such as integrating pegRNA/target library and then Cas9, the same Lipofectamine 3000 transfection protocol with Tol2 transposase plasmid was performed each time, and >1 week of appropriate drug selection was performed after each transfection.

### Deep sequencing, library preparation

Genomic DNA was extracted from harvested cells with the PureLink Genomic DNA Purification Mini Kit (Invitrogen). For library experiments, sequences including the peptide and the prime editing site were PCR amplified using Q5® High-Fidelity DNA Polymerase (NEB) and primers as specified (Supplementary Table 6). For each replicate, the first PCR included a total of 10-20 μg of genomic DNA. To determine the number of cycles required to complete the exponential phase of amplification we first performed qPCR, followed by PCR using primers that included both Illumina adaptor and barcode sequences (Supplementary Table 6). For measuring PE2 efficiencies at endogenous sites, the independent first PCR was performed in a 200ul reaction volume that contained 1000ng of the initial genomic DNA template per sample. The second PCR to attach the Illumina adaptor and barcode sequences was then performed using purified product from the first PCR. After bead purification, pooled samples were sequenced using NextSeq (Illumina).

### Library data processing

Designed library peptides were identified in sequenced reads by exact string matching to the first eight nucleotides of the peptide sequence, which were unique across the library. Sequenced target sites were aligned to the designed reference using Needleman-Wunsch with match score 1, mismatch cost −1, gap open cost −5, and gap extend cost 0. Reads with mean PHRED quality score below 30 were filtered. Mismatches at nucleotides with less than PHRED quality score 30 were filtered. Indels with less than three matching nucleotides on both sides with at least PHRED quality score 30 were filtered.

### Identifying peptide hits

We excluded peptides with less than 100 reads in any experiment. We used a beta-binomial model to infer peptide editing effects from replicate data while adjusting for sampling noise from limited sequencing reads. We model a peptide *i* with parameters α_*i*_, β_*i*_ used to sample a peptide effect *p*_*ij*_ *∼ Beta*(α_*i*_, β_*i*_) for an experiment or batch *j*. Samples from experiment or batch *j* are taken for sequencing, yielding a binomial distribution over the number of edited reads *y*_*ij*_ *∼ Bin*(*n*_*j*_, *p*_*ij*_) for read depth *n*_*j*_. Given *k* samples of *y*_*ij*_ over *k* biological replicates or batches, we infer the maximum likelihood estimate (MLE) of α_*i*_, β_*i*_ for peptide *i*. As our beta-binomial model is conjugate, the MLE of α_*i*_, β_*i*_ can be found analytically by solving the system of equations:

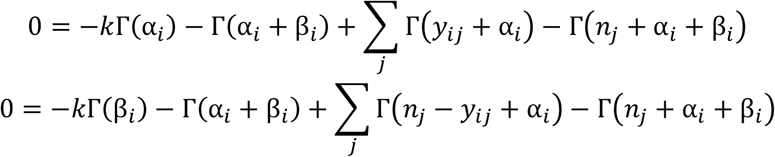

Where Γ() is the Gamma function. We solved this using Sympy^1^. When solutions could not be found due to numerical instability, we used a fast approximation that solves the MLE of α<_*i*_, β_*i*_ by matching the observed mean and variance, motivated by viewing the beta-binomial distribution as an overdispersed binomial distribution. The additional variance over a binomial distribution is related to the sum α_*i*_ + β_*i*_.

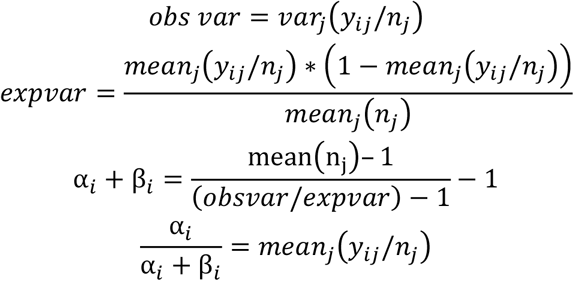

To interpret α_*i*_, β_*i*_, we convert them into the mean 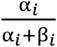 and variance 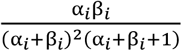 of a Beta distribution.

We selected peptides for follow-up evaluation using several metrics. To increase confidence in hits, we preferred peptides present in higher numbers of replicates. We prioritized peptides based on the probability of observing a higher edited read count under its inferred peptide effect parameters compared to edited read counts sampled from inferred control peptide effect parameters under our beta-binomial model, which prefers higher MLE mean and lower MLE variance. We also selected peptides with high MLE mean effect even if their variance was high.

### 100 target site library design

An oligonucleotide pool containing 100 target sequences was synthesized by IDT. Each oligonucleotide contained the following elements 3’ to 5’: 19nt PE1 stub, 4nt barcode, ∼40nt variable target, 6nt poly A terminator, ∼30nt PBS/template, 86nt hairpin, 20nt spacer, 20nt U6 stub. The barcode stuffer allowed individual pegRNA and target sequence pairs to be identified after deep sequencing. To test the effect of PBS and RT template length on PE2 efficiency, we prepared pegRNAs with 8 different combinations of edit types. Types of mutations:

- 3 × 1-nt substitution
  - PAM NGG-->NCG
  - PAM NGG-->NGT
  - Seed 1 transversion—nt nearest PAM, AàT, TàA, CàG, GàC
- 3 x >1-nt substitution

- PAM NGG-->NCT
- Seed 2-3 transversion (AàT, TàA, CàG, GàC) + PAM NGGàNTC [discontinuous, maintain 2 intervening nt]
- 6-nt PAM + seed change to GAATTC
- 1 × 1-nt ins

- PAM NGG-->NGTG
- 1 × 1-nt del
  - PAM NGG-->not NGG

- delete 1st G unless G after PAM
- Delete seed 1 unless 1st base of PAM is identical

To design a library of 100 pegRNA-target pairs we used 96 pegRNA-target pairs from Kim et al high-throughput library data that vary in prime editing efficiency. In their library, for all sites with DeepSpCas9 score >20, average PE efficiency is 11%, SD=9%. We chose 4 target sites 0-1 SD below average (1%, 3%, 5%, 8% PE), 4 sites around average (11%, 14%, 17%, 20%), 2 sites ∼1 SD above average (30%, 40%), 2 near top (50%, 60%). Our library also includes 4 substitution mutations from Anzalone et al that showed highest prime editing activity. Oligo library was cloned into pLenti-sgRNA-FE vector using NEBuilder HiFi DNA Assembly Kit (NEB). Assembled plasmids were purified by isopropanol precipitation with GlycoBlue Coprecipitant (Thermo Fisher) and reconstituted in TE and transformed into Endura™ Electrocompetent Cells (Lucigen). After library diversity was verified, library mastermix was used to produce lentivirus.

### Production of lentivirus and cellular infection

HEK293FT cells (15 × 10^6) were seeded on 150-mm cell culture dishes containing DMEM. The next day, cells were transfected with pCMV-VSV-G (Addgene #8454), pRSV-Rev (Addgene #12253), pMDLg/pRRE (Addgene #12251) and library, at a ratio of 1:2:3:4, using TransIT®-Lenti Transfection Reagent (Mirus Bio). At 8h after transfection, cells were refreshed with maintaining medium. At 24h and 48h after transfection, the lentivirus-containing supernatant was collected, filtered through a 0.45-μm pore filter (Corning), concentrated using Lenti-X™ Concentrator (TakaraBio), aliquoted and stored at −80°C.

In preparation for lentivirus transduction, cells (U2OS, HCT-116, mESC, HEK293FT) were seeded on 100-mm dishes (at a density of 2 × 10^5, 6.5 × 10^4, 6.5 × 10^4, 1 × 10^5 cells per cm^2) and concentrated lenti was added to the media. The cells were then incubated overnight, after which cells were refreshed with maintaining medium before adding blasticidin at 48h and keeping it for minimum of next 5 d to remove untransduced cells. To preserve its diversity, the cell library was maintained at a count of at least 1 × 10^7 cells throughout the study.

### Measurement of PE2 efficiencies at endogenous sites

To validate the results of the high-throughput experiments, 6 individual pegRNA-encoding plasmids targeting endogenous NF2 locus were constructed and used to produce lentiviral particles. In preparation for transfection, HEK293T and U2OS cells were seeded on 10 cm plates at a density of 4*10^4^ cells per cm^2^ and transduced with a lentivirus carrying pegRNA-encoding plasmid. After cells were selected for successful lentiviral integration, they were transfected using Lipofectamine 3000 with plasmid encoding IN-PE2 or PE2-control and equimolar amounts of Tol2 transposase plasmid, according to the manufacturer’s instructions. After a week of selection for successful integration of constructs, cells were harvested for gDNA extraction followed by library preparation for NGS. Primers used to sequence NF2 locus are listed in Supplementary table 6.

## Code availability

Custom code used to process and analyze peptide library data are available at https://github.com/maxwshen/prime-peptide.

## Notes

### Competing Interest Statement

The authors have declared no competing interest.

## References

1. Anzalone, A. V. et al. Search-and-replace genome editing without double-strand breaks or donor DNA. Nature 576, 149–157 (2019).

2. Anzalone, A. V., Koblan, L. W. & Liu, D. R. Genome editing with CRISPR-Cas nucleases, base editors, transposases and prime editors. Nat. Biotechnol. 38, 824–844 (2020).

3. Kim, H. K. et al. Predicting the efficiency of prime editing guide RNAs in human cells. Nat. Biotechnol. 1–9 (2020) doi:10.1038/s41587-020-0677-y.

4. Kim, H. K. et al. SpCas9 activity prediction by DeepSpCas9, a deep learning–based model with high generalization performance. Sci. Adv. 5, eaax9249 (2019).

5. Lin, Q. et al. High-efficiency prime editing with optimized, paired pegRNAs in plants. Nat. Biotechnol. (2021) doi:10.1038/s41587-021-00868-w.

6. Zhang, X. et al. Increasing the efficiency and targeting range of cytidine base editors through fusion of a single-stranded DNA-binding protein domain. Nat. Cell Biol. 22, 740–750 (2020).

7. Komor, A. C., Kim, Y. B., Packer, M. S., Zuris, J. A. & Liu, D. R. Programmable editing of a target base in genomic DNA without double-stranded DNA cleavage. Nature 533, 420–424 (2016).

8. Pickar-Oliver, A. & Gersbach, C. A. The next generation of CRISPR–Cas technologies and applications. Nat. Rev. Mol. Cell Biol. 20, 490–507 (2019).

9. Owusu, M. et al. Mapping the Human Kinome in Response to DNA Damage. Cell Rep. 26, 555–563.e6 (2019).

10. Wood, R. D., Mitchell, M. & Lindahl, T. Human DNA repair genes, 2005. Mutat. Res. 577, 275–283 (2005).

11. Sherwood, R. I. et al. Discovery of directional and nondirectional pioneer transcription factors by modeling DNase profile magnitude and shape. Nat Biotechnol 32, 171–8 (2014).

12. Lin, L. et al. Comprehensive Mapping of Key Regulatory Networks that Drive Oncogene Expression. Cell Rep. 33, 108426 (2020).

13. Trojan, J. et al. Functional analysis of hMLH1 variants and HNPCC-related mutations using a human expression system. Gastroenterology 122, 211–219 (2002).

14. Papadopoulos, N. et al. Mutation of a mutL homolog in hereditary colon cancer. Science 263, 1625 (1994).

15. Xu, G. J. et al. Systematic autoantigen analysis identifies a distinct subtype of scleroderma with coincident cancer. Proc. Natl. Acad. Sci. 113, E7526 (2016).

16. Knijnenburg, T. A. et al. Genomic and Molecular Landscape of DNA Damage Repair Deficiency across The Cancer Genome Atlas. Cell Rep. 23, 239–254.e6 (2018).

17. Friedberg, E. C. et al. DNA Repair and Mutagenesis. (ASM Press, 2005). doi:10.1128/9781555816704.

18. Shen, M. W. et al. Predictable and precise template-free CRISPR editing of pathogenic variants. Nature 563, 646–651 (2018).

