## Supplemental File for "Peptide fusion improves prime editing efficiency"

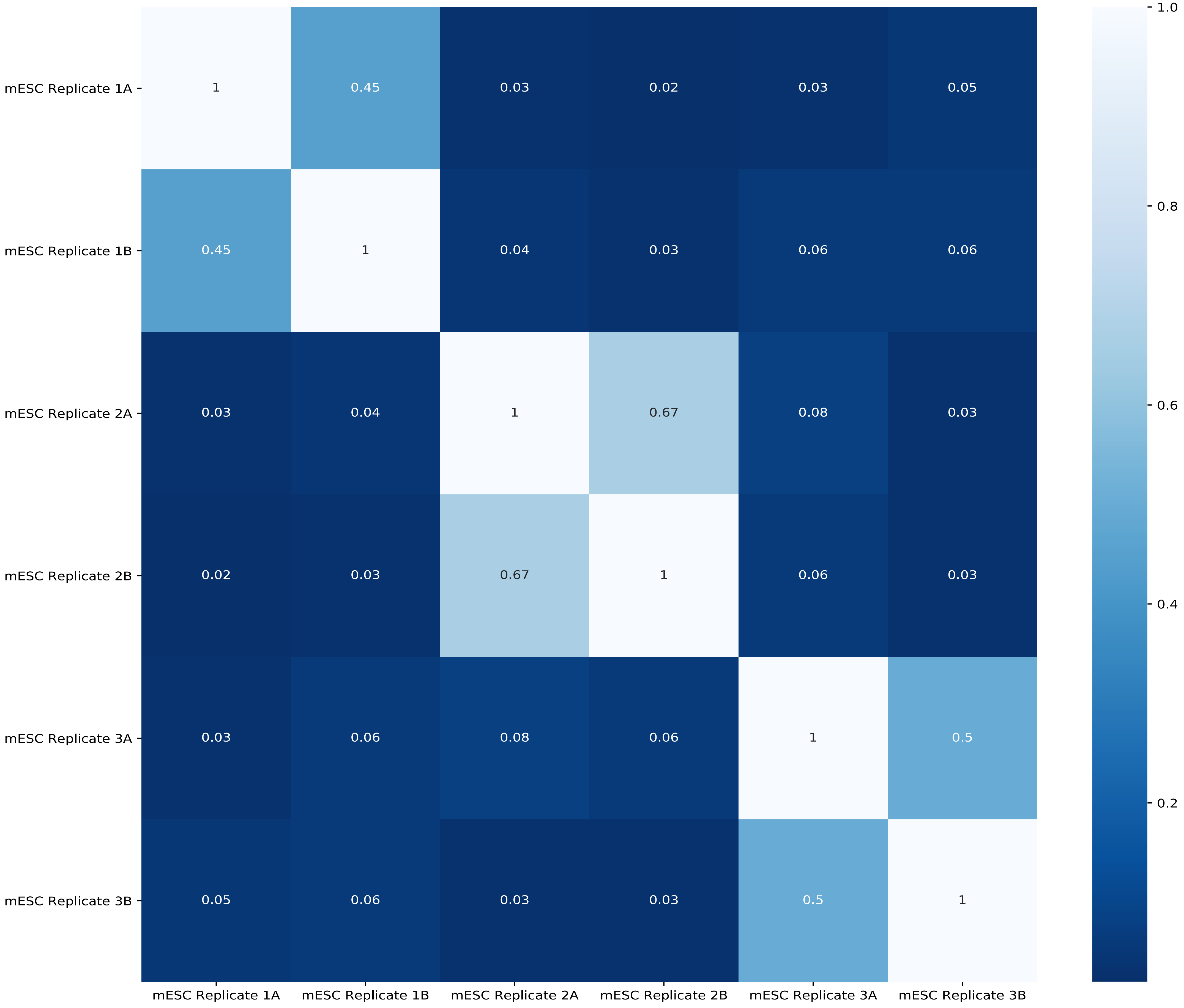


**Supplementary Figure 1. Replicate consistency in 12,000-peptide PepSEq screens.**
(a) Comparison of prime editing efficiency in 12,000-peptide PepSEq screens for all pairs of replicates with independent and shared peptide-PE2 integration. Technical replicates were combined and biological replicates were used for analysis.

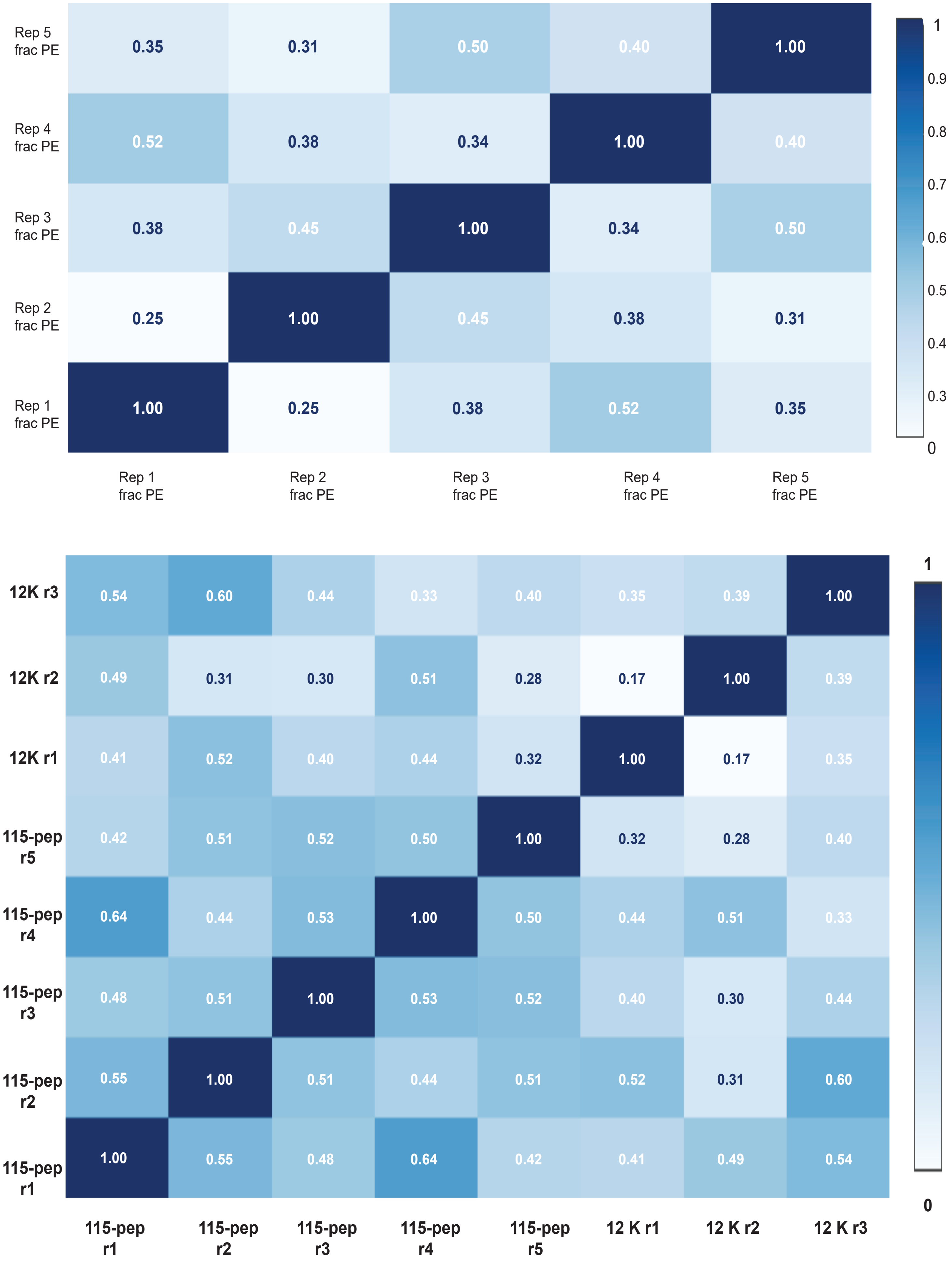


**Supplementary Figure 2: A 115-peptide PepSEq screen reveals peptides that induce replicate-consistent increases in prime editing efficiency.**

(a) Correlation table of prime editing efficiency in five biological replicate 115-peptide PepSEq screens.

(b) Correlation table of prime editing efficiency of 45 strongest prime editing-enhancing peptides and 10 control peptides in five biological replicate 115-peptide PepSEq screens and three biological replicate 12,000-peptide screens.

| | Ctrl-norm frac PE mESC r1 | Ctrl-norm frac PE mESC r2 | Ctrl-norm frac PE mESC r3 | Ctrl-norm frac PE mESC r4 | Ctrl-norm frac PE mESC r5 | Ctrl-norm frac PE mESC r6 | Ctrl-norm frac PE mESC r7 | Ctrl-norm frac PE mESC r8 | Ctrl-norm frac PE mESC r9 | Ctrl-norm frac PE mESC r10 | Ctrl-norm frac PE HCT116 r1 | Ctrl-norm frac PE HCT116 r2 |
|---|---|---|---|---|---|---|---|---|---|---|---|---|
| Ctrl-norm frac PE HCT116 r2 | 0.31 | 0.25 | 0.21 | 0.14 | 0.29 | 0.23 | 0.33 | 0.22 | 0.10 | 0.29 | 0.47 | 1.00 |
| Ctrl-norm frac PE HCT116 r1 | 0.36 | 0.35 | 0.34 | 0.34 | 0.42 | 0.26 | 0.42 | 0.41 | 0.42 | 0.47 | 1.00 | 0.47 |
| Ctrl-norm frac PE mESC r10 | 0.60 | 0.63 | 0.62 | 0.58 | 0.65 | 0.65 | 0.60 | 0.57 | 0.49 | 1.00 | 0.47 | 0.29 |
| Ctrl-norm frac PE mESC r9 | 0.55 | 0.62 | 0.51 | 0.61 | 0.54 | 0.43 | 0.59 | 0.68 | 1.00 | 0.49 | 0.42 | 0.10 |
| Ctrl-norm frac PE mESC r8 | 0.65 | 0.70 | 0.68 | 0.69 | 0.63 | 0.58 | 0.67 | 1.00 | 0.68 | 0.57 | 0.41 | 0.22 |
| Ctrl-norm frac PE mESC r7 | 0.75 | 0.68 | 0.62 | 0.69 | 0.64 | 0.57 | 1.00 | 0.67 | 0.59 | 0.60 | 0.42 | 0.33 |
| Ctrl-norm frac PE mESC r6 | 0.48 | 0.54 | 0.63 | 0.55 | 0.63 | 1.00 | 0.57 | 0.58 | 0.43 | 0.65 | 0.26 | 0.23 |
| Ctrl-norm frac PE mESC r5 | 0.66 | 0.70 | 0.60 | 0.67 | 1.00 | 0.63 | 0.64 | 0.63 | 0.54 | 0.65 | 0.42 | 0.29 |
| Ctrl-norm frac PE mESC r4 | 0.61 | 0.65 | 0.67 | 1.00 | 0.67 | 0.55 | 0.69 | 0.69 | 0.61 | 0.58 | 0.34 | 0.14 |
| Ctrl-norm frac PE mESC r3 | 0.57 | 0.62 | 1.00 | 0.67 | 0.60 | 0.63 | 0.62 | 0.68 | 0.51 | 0.62 | 0.34 | 0.21 |
| Ctrl-norm frac PE mESC r2 | 0.62 | 1.00 | 0.62 | 0.65 | 0.70 | 0.54 | 0.68 | 0.70 | 0.62 | 0.62 | 0.35 | 0.25 |
| Ctrl-norm frac PE mESC r1 | 1.00 | 0.62 | 0.57 | 0.61 | 0.66 | 0.48 | 0.75 | 0.65 | 0.55 | 0.60 | 0.36 | 0.31 |

0

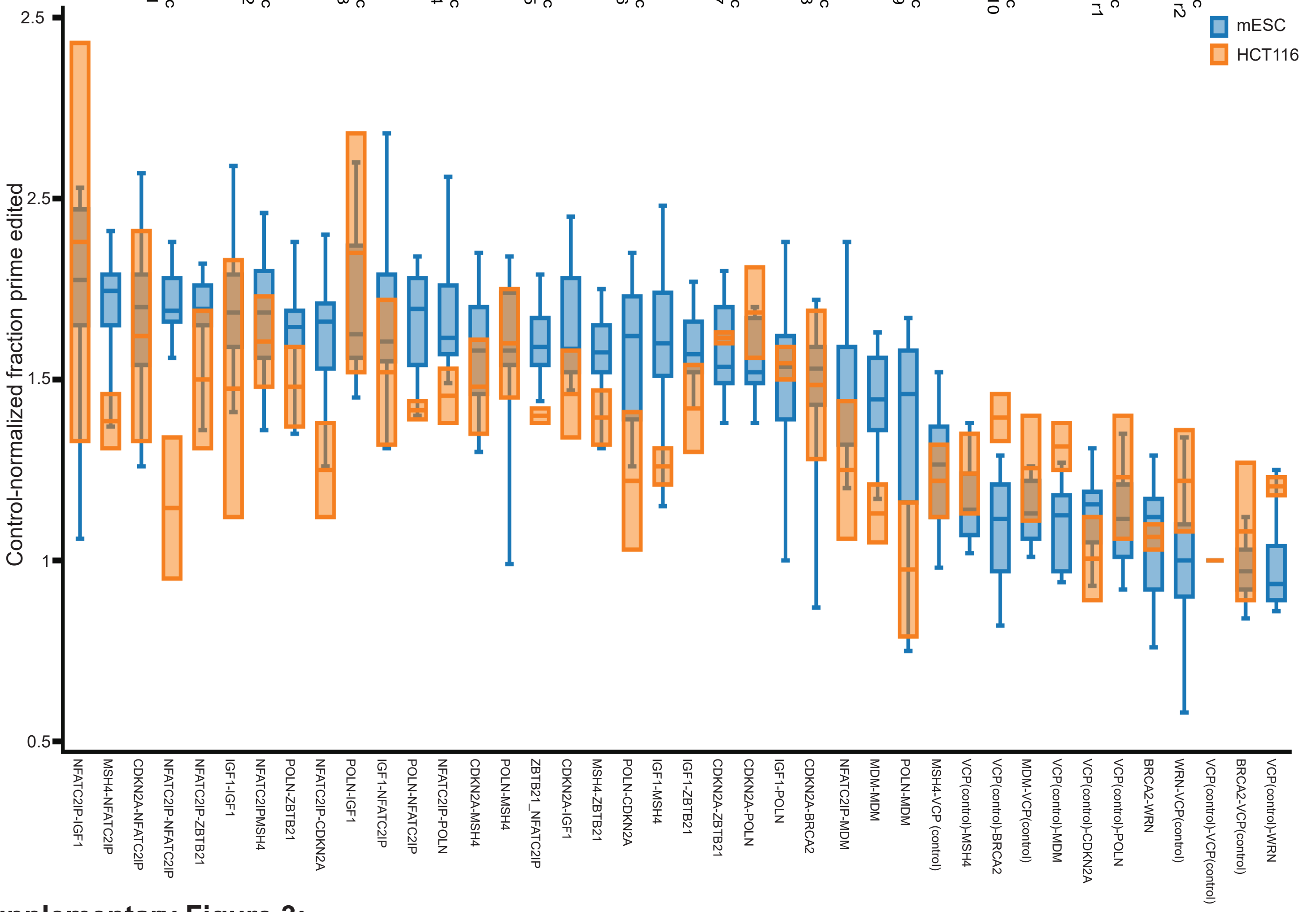


**Supplementary Figure 3:**

**A dual-peptide PepSEq screen reveals linear contribution of prime editing-enhancing peptides.**

(a) Correlation table of prime editing efficiency in 10 biological replicate dual-peptide PepSEq screens in mESC and two biological replicates in HCT-116.

(b) Comparison of control-normalized prime edited fraction for 100 dual-peptide pairs in mESC and HCT-116.

(c) Comparison of inferred effect of each peptide from dual-peptide linear model vs. observed peptide effects from 115-peptide screen.

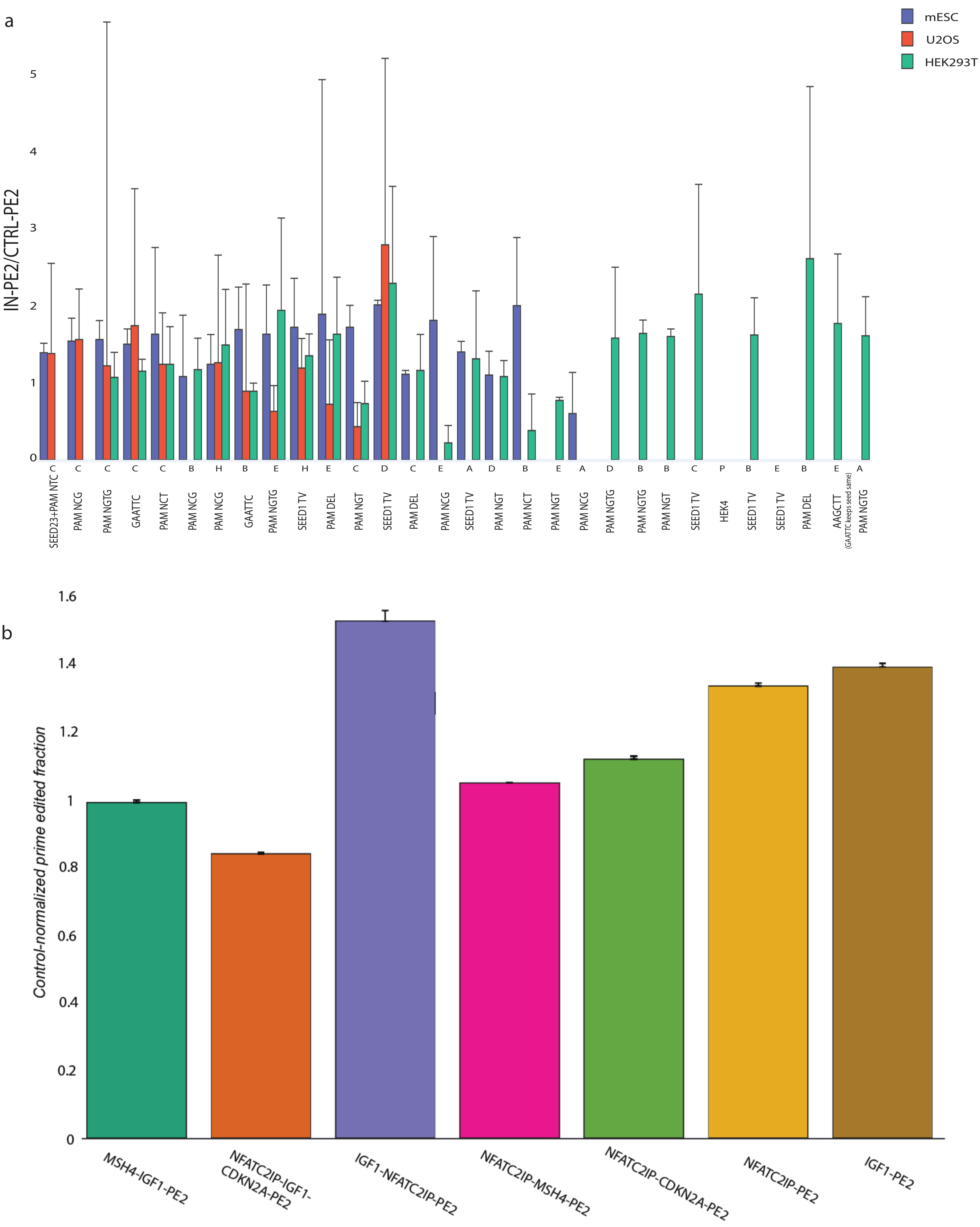


**Supplementary Figure 4: Peptide-PE2 editing in a 100-member pegRNA-target library**

(a) Comparison of control-normalized prime edited fraction for IN-PE2 and CTRL-PE2 at up to 30 target sites in three cell lines.

(b) Median control-normalized prime edited fraction for seven single, double, and triple-peptide-PE2s as compared with CTRL-PE2 at up to 30 target sites in mESC and U2OS.

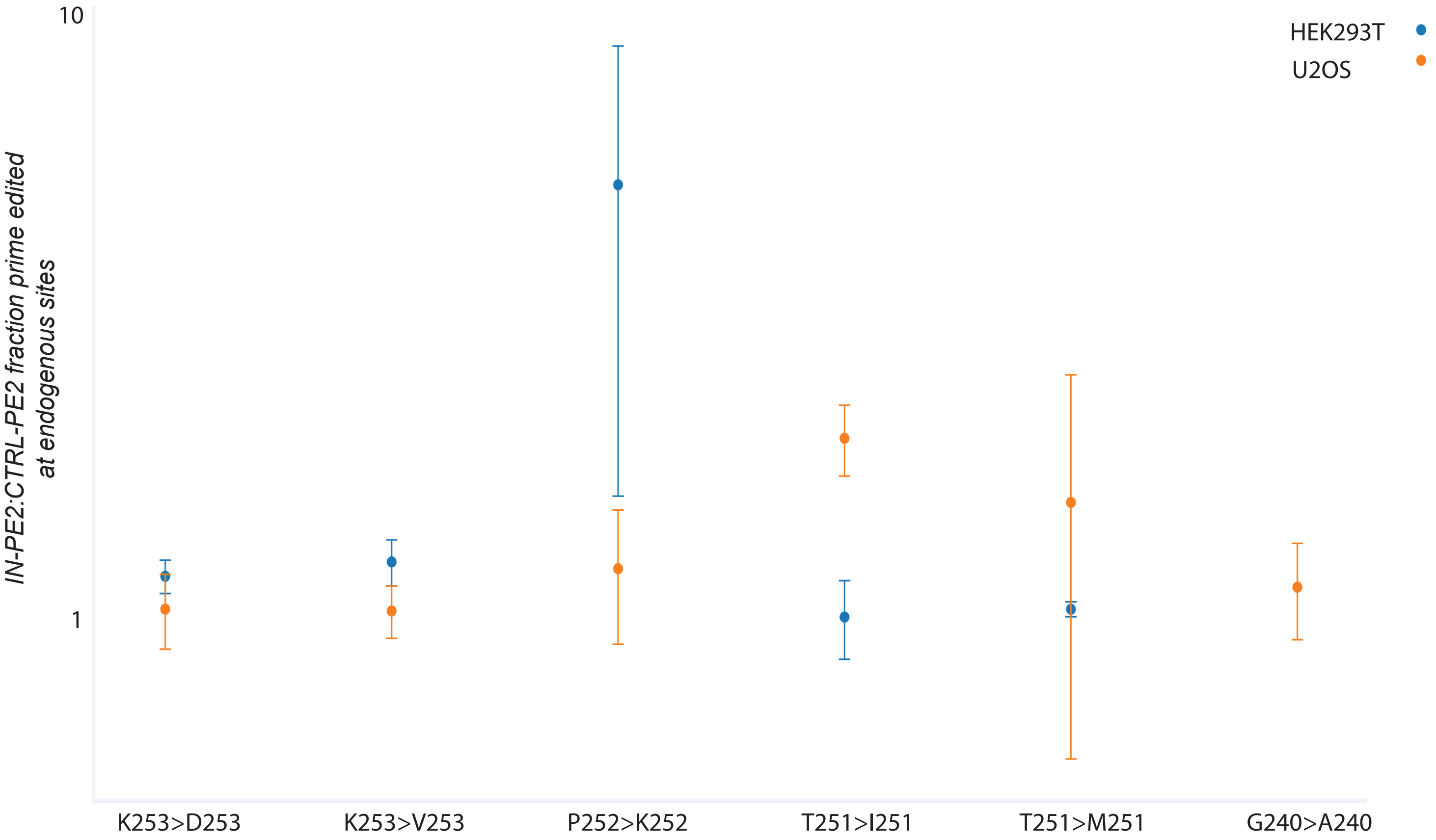


**Supplementary Figure 5: IN-PE2 editing at endogenous human loci.**

a) Control-normalized prime edited fraction at NF2 endogenous human loci in HEK293T and U2OS cells

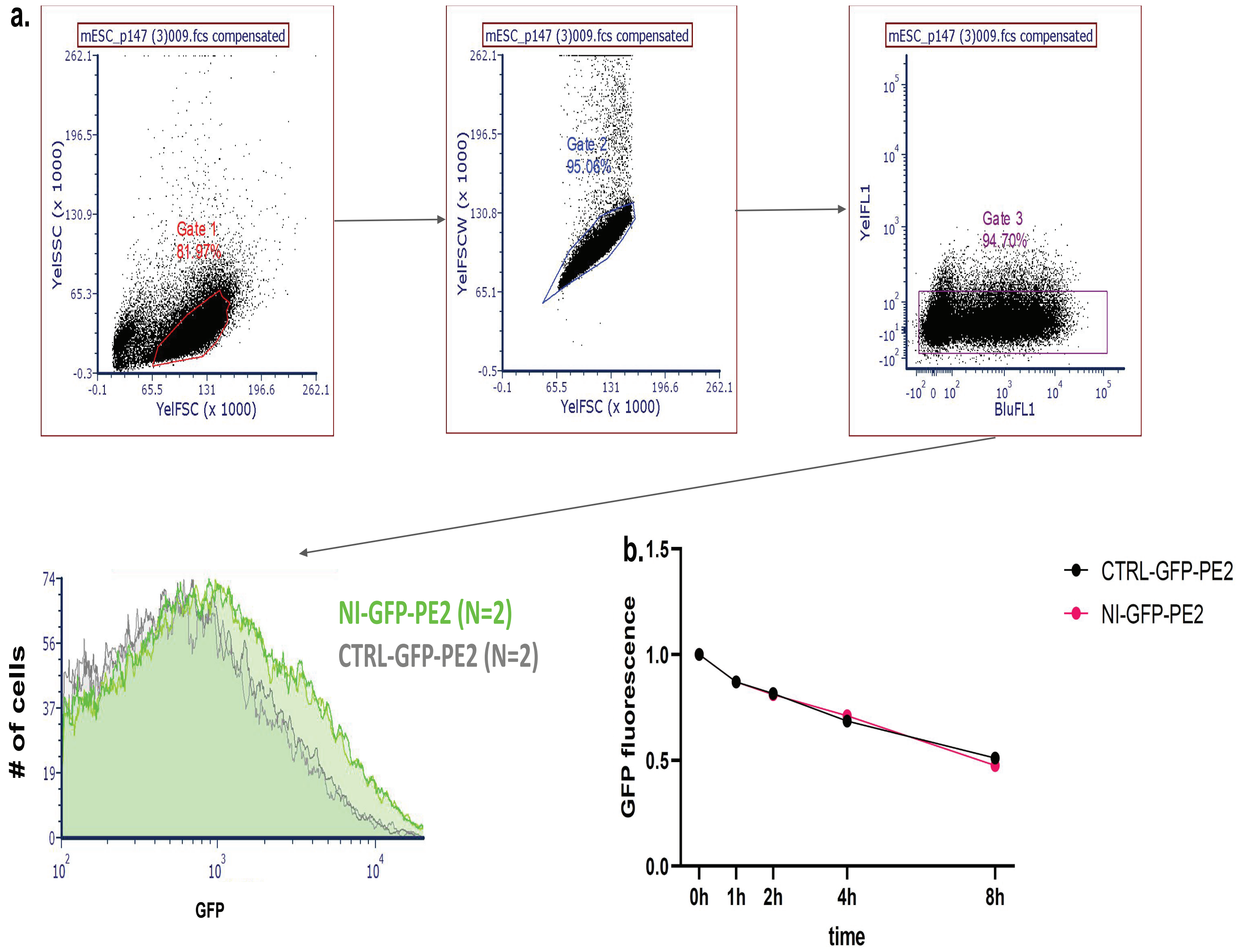


**Supplementary Figure 6: IN-PE2 is expressed at an increased level as compared to CTRL-PE2.**

(a) Full gating for flow cytometric analysis of IN-GFP-PE2 and CTRL-GFP-PE2.

(b) IN-GFP-PE2 and CTRL-GFP-PE2 fluorescence during an 8-hour cycloheximide timecourse, normalized to fluorescence in untreated cells.

**MSH4-IGF1-NFATC2IP-PE2 ORF**

QVVSASTCPGTSGAAGDRSSSSSSLPCPAPNSRPAQGSYFGNKRAYAENTVASNFTFGASSSSARDTNYPQTLKTPLST
GNPQRS GSTGGTGS
REGMNKPEGYGEEERRAPQEGIVDECCFRECDLRRLEMYCAPLKPAKEAREVRAQRHEDMPKEQKYQPPEENKNEKE
QRRKGEEF GSTGGTGS
AEPVGKRGRWSGGSGAGRGGRGGWGGRGRRPRAQRSPSRGTLDVVSVDLVTDSDEEILEVATARGAADEVEVEPPE
PPGPVASRD SSGGSSGGSSGSETPGTSESATPESSGGSSGGSCT
DKKYSIGLDIGTNSVGWAVITDEYKVPSKKFKVLGNTDRHSIKKNLIGALLFDSGETAEATRLKRTARRRYTRRKNRICYLQ
EIFSNEMAKVDDSFFHRLEESFLVEEDKKHERHPIFGNIVDEVAYHEKYPTIYHLRKKLVDSTDKADLRLIYLALAHMIKFR
GHFLIEGDLNPDNSDVDKLFIQLVQTYNQLFEENPINASGVDAKAILSARLSKSRRLENLIAQLPGEKKNGLFGNLIALSLG
LTPNFKSNFDLAEDAKLQLSKDTYDDDLDNLLAQIGDQYADLFLAAKNLSDAILLSDILRVNTEITKAPLSASMIKRYDEHH
QDLTLLKALVRQQLPEKYKEIFFDQSKNGYAGYIDGGASQEEFYKFIKPILEKMDGTEELLVKLNREDLLRKQRTFDNGSIP
HQIHLGELHAILRRQEDFYPFLKDNREKIEKILTFRIPYYVGPLARGNSRFAWMTRKSEETITPWNFEEVVDKGASAQSFI
ERMTNFDKNLPNEKVLPKHSLLYEYFTVYNELTKVKYVTEGMRKPAFLSGEQKKAIVDLLFKTNRKVTVKQLKEDYFKKIE
CFDSVEISGVEDRFNASLGTYHDLLKIIKDKDFLDNEENEDILEDIVLTLTLFEDREMIEERLKTYAHLFDDKVMKQLKRRR
YTGWGRLSRKLINGIRDKQSGKTILDFLKSDGFANRNFMQLIHDDSLTFKEDIQKAQVSGQGDSLHEHIANLAGSPAIKK
GILQTVKVVDELVKVMGRHKPENIVIEMARENQTTQKGQKNSRERMKRIEEGIKELGSQILKEHPVENTQLQNEKLYLY
YLQNGRDMYVDQELDINRLSDYDVDAIVPQSFLKDDSIDNKVLTRSDKNRGKSDNVPSEEVVKKMKNYWRQLLNAKLI
TQRKFDNLTKAERGGLSELDKAGFIKRQLVETRQITKHVAQILDSRMNTKYDENDKLIREVKVITLKSKLVSDFRKDFQFY
KVREINNYHHAHDAYLNAVVGTALIKKYPKLESEFVYGDYKVYDVRKMIAKSEQEIGKATAKYFFYSNIMNFFKTEITLAN
GEIRKRPLIETNGETGEIVWDKGRDFATVRKVLSMPQVNIVKKTEVQTGGFSKESILPKRNSDKLIARKKDWDPKKYGGF
DSPTVAYSVLVVAKVEKGKSKKLKSVKELLGITIMERSSFEKNPIDFLEAKGYKEVKKDLIIKLPKYSLFELENGRKRMLASA
GELQKGNELALPSKYVNFLYLASHYEKLKGSPEDNEQKQLFVEQHKHYLDEIIEQISEFSKRVILADANLDKVLSAYNKHR
DKPIREQAENIIHLFTLTNLGAPAAFKYFDTTIDRKRYTSTKEVLDATLIHQSITGLYETRIDLSQLGGD
TLNIEDEYRLHETSKEPDVSLGSTWLSDFPQAWAETGGMGLAVRQAPLIIPLKATSTPVSIKQYPMSQEARLGIKPHIQR
LLDQGILVPCQSPWNTPLLPVKKPGTNDYRPVQDLREVNKRVEDIHPTVPNPYNLLSGLPPSHQWYTVLDLKDAFFCLR
LHPTSQPLFAFEWRDPEMGISGQLTWTRLPQGFKNSPTLFNEALHRDLADFRIQHPDLILLQYVDDLLLAATSELDCQQ
GTRALLQTLGNLGYRASAKKAQICQKQVKYLGYLLKEGQRWLTEARKETVMGQPTPKTPRQLREFLGKAGFCRLFIPGF
AEMAAPLYPLTKPGTLFNWGPDQQKAYQEIKQALLTAPALGLPDLTKPFELFVDEKQGYAKGVLTQKLGPWRRPVAYL
SKKLDPVAAGWPPCLRMVAAIAVLTKDAGKLTMGQPLVILAPHAVEALVKQPPDRWLSNARMTHYQALLLDTDRVQ
FGPVVALNPATLLPLPEEGLQHNCLDILAEAHGTRPDLTDQPLPDADHTWYTDGSSLLQEGQRKAGAAVTTETEVIWA
KALPAGTSAQRAELIALTQALKMAEGKKLNVYTDSRYAFATAHIHGEIYRRRGWLTSEGKEIKNKDEILALLKALFLPKRLS
IIHCPGHQKGHSAEARGNRMADQAARKAAITETPDTSTLLIENSSP

**NFATC2IP-IGF1-CDKN2A-PE2 ORF**

AEPVGKRGRWSGGSGAGRGGRGGWGGRGRRPRAQRSPSRGTLDVVSVDLVTDSDEEILEVATARGAADEVEVEPPE
PPGPVASRD GSTGGTGS
REGMNKPEGYGEEERRAPQEGIVDECCFRECDLRRLEMYCAPLKPAKEAREVRAQRHEDMPKEQKYQPPEENKNEKE
QRRKGEEF GSTGGTGS
EPAAGSSMEPSADWLATAAARGRVEEVRALLEAGALPNAPNSYGRRPIQVMMMGSARVAELLLLHGAEPNCADPAT
LTRPVHDAA SSGGSSGGSSGSETPGTSESATPESSGGSSGGSCT
DKKYSIGLDIGTNSVGWAVITDEYKVPSKKFKVLGNTDRHSIKKNLIGALLFDSGETAEATRLKRTARRRYTRRKNRICYLQ
EIFSNEMAKVDDSFFHRLEESFLVEEDKKHERHPIFGNIVDEVAYHEKYPTIYHLRKKLVDSTDKADLRLIYLALAHMIKFR
GHFLIEGDLNPDNSDVDKLFIQLVQTYNQLFEENPINASGVDAKAILSARLSKSRRLENLIAQLPGEKKNGLFGNLIALSLG
LTPNFKSNFDLAEDAKLQLSKDTYDDDLDNLLAQIGDQYADLFLAAKNLSDAILLSDILRVNTEITKAPLSASMIKRYDEHH
QDLTLLKALVRQQLPEKYKEIFFDQSKNGYAGYIDGGASQEEFYKFIKPILEKMDGTEELLVKLNREDLLRKQRTFDNGSIP
HQIHLGELHAILRRQEDFYPFLKDNREKIEKILTFRIPYYVGPLARGNSRFAWMTRKSEETITPWNFEEVVDKGASAQSFI
ERMTNFDKNLPNEKVLPKHSLLYEYFTVYNELTKVKYVTEGMRKPAFLSGEQKKAIVDLLFKTNRKVTVKQLKEDYFKKIE

CFDSVEISGVEDRFNASLGTYHDLLKIIKDKDFLDNEENEDILEDIVLTLTLFEDREMIEERLKTYAHLFDDKVMKQLKRRR
YTGWGRLSRKLINGIRDKQSGKTILDFLKSDGFANRNFMQLIHDDSLTFKEDIQKAQVSGQGDSLHEHIANLAGSPAIKK
GILQTVKVVDELVKVMGRHKPENIVIEMARENQTTQKGQKNSRERMKRIEEGIKELGSQILKEHPVENTQLQNEKLYLY
YLQNGRDMYVDQELDINRLSDYDVDAIVPQSFLKDDSIDNKVLTRSDKNRGKSDNVPSEEVVKKMKNYWRQLLNAKLI
TQRKFDNLTKAERGGLSELDKAGFIKRQLVETRQITKHVAQILDSRMNTKYDENDKLIREVKVITLKSKLVSDFRKDFQFY
KVREINNYHHAHDAYLNAVVGTALIKKYPKLESEFVYGDYKVYDVRKMIAKSEQEIGKATAKYFFYSNIMNFFKTEITLAN
GEIRKRPLIETNGETGEIVWDKGRDFATVRKVLSMPQVNIVKKTEVQTGGFSKESILPKRNSDKLIARKKDWDPKKYGGF
DSPTVAYSVLVVAKVEKGKSKKLKSVKELLGITIMERSSFEKNPIDFLEAKGYKEVKKDLIIKLPKYSLFELENGRKRMLASA
GELQKGNELALPSKYVNFLYLASHYEKLKGSPEDNEQKQLFVEQHKHYLDEIIEQISEFSKRVILADANLDKVLSAYNKHR
DKPIREQAENIIHLFTLTNLGAPAAFKYFDTTIDRKRYTSTKEVLDATLIHQSITGLYETRIDLSQLGGD
TLNIEDEYRLHETSKEPDVSLGSTWLSDFPQAWAETGGMGLAVRQAPLIIPLKATSTPVSIKQYPMSQEARLGIKPHIQR
LLDQGILVPCQSPWNTPLLPVKKPGTNDYRPVQDLREVNKRVEDIHPTVPNPYNLLSGLPPSHQWYTVLDLKDAFFCLR
LHPTSQPLFAFEWRDPEMGISGQLTWTRLPQGFKNSPTLFNEALHRDLADFRIQHPDLILLQYVDDLLLAATSELDCQQ
GTRALLQTLGNLGYRASAKKAQICQKQVKYLGYLLKEGQRWLTEARKETVMGQPTPKTPRQLREFLGKAGFCRLFIPGF
AEMAAPLYPLTKPGTLFNWGPDQQKAYQEIKQALLTAPALGLPDLTKPFELFVDEKQGYAKGVLTQKLGPWRRPVAYL
SKKLDPVAAGWPPCLRMVAAIAVLTKDAGKLTMGQPLVILAPHAVEALVKQPPDRWLSNARMTHYQALLLDTDRVQ
FGPVVALNPATLLPLPEEGLQHNCLDILAEAHGTRPDLTDQPLPDADHTWYTDGSSLLQEGQRKAGAAVTTETEVIWA
KALPAGTSAQRAELIALTQALKMAEGKKLNVYTDSRYAFATAHIHGEIYRRRGWLTSEGKEIKNKDEILALLKALFLPKRLS
IIHCPGHQKGHSAEARGNRMADQAARKAAITETPDTSTLLIENSSP

**IGF1-NFATC2IP-PE2 ORF**

REGMNKPEGYGEEERRAPQEGIVDECCFRECDLRRLEMYCAPLKPAKEAREVRAQRHEDMPKEQKYQPPEENKNEKE
QRRKGEEF GSTGGTGS
AEPVGKRGRWSGGSGAGRGGRGGWGGRGRRPRAQRSPSRGTLDVVSVDLVTDSDEEILEVATARGAADEVEVEPPE
PPGPVASRD SSGGSSGGSSGSETPGTSESATPESSGGSSGGSCT
DKKYSIGLDIGTNSVGWAVITDEYKVPSKKFKVLGNTDRHSIKKNLIGALLFDSGETAEATRLKRTARRRYTRRKNRICYLQ
EIFSNEMAKVDDSFFHRLEESFLVEEDKKHERHPIFGNIVDEVAYHEKYPTIYHLRKKLVDSTDKADLRLIYLALAHMIKFR
GHFLIEGDLNPDNSDVDKLFIQLVQTYNQLFEENPINASGVDAKAILSARLSKSRRLENLIAQLPGEKKNGLFGNLIALSLG
LTPNFKSNFDLAEDAKLQLSKDTYDDDLDNLLAQIGDQYADLFLAAKNLSDAILLSDILRVNTEITKAPLSASMIKRYDEHH
QDLTLLKALVRQQLPEKYKEIFFDQSKNGYAGYIDGGASQEEFYKFIKPILEKMDGTEELLVKLNREDLLRKQRTFDNGSIP
HQIHLGELHAILRRQEDFYPFLKDNREKIEKILTFRIPYYVGPLARGNSRFAWMTRKSEETITPWNFEEVVDKGASAQSFI
ERMTNFDKNLPNEKVLPKHSLLYEYFTVYNELTKVKYVTEGMRKPAFLSGEQKKAIVDLLFKTNRKVTVKQLKEDYFKKIE
CFDSVEISGVEDRFNASLGTYHDLLKIIKDKDFLDNEENEDILEDIVLTLTLFEDREMIEERLKTYAHLFDDKVMKQLKRRR
YTGWGRLSRKLINGIRDKQSGKTILDFLKSDGFANRNFMQLIHDDSLTFKEDIQKAQVSGQGDSLHEHIANLAGSPAIKK
GILQTVKVVDELVKVMGRHKPENIVIEMARENQTTQKGQKNSRERMKRIEEGIKELGSQILKEHPVENTQLQNEKLYLY
YLQNGRDMYVDQELDINRLSDYDVDAIVPQSFLKDDSIDNKVLTRSDKNRGKSDNVPSEEVVKKMKNYWRQLLNAKLI
TQRKFDNLTKAERGGLSELDKAGFIKRQLVETRQITKHVAQILDSRMNTKYDENDKLIREVKVITLKSKLVSDFRKDFQFY
KVREINNYHHAHDAYLNAVVGTALIKKYPKLESEFVYGDYKVYDVRKMIAKSEQEIGKATAKYFFYSNIMNFFKTEITLAN
GEIRKRPLIETNGETGEIVWDKGRDFATVRKVLSMPQVNIVKKTEVQTGGFSKESILPKRNSDKLIARKKDWDPKKYGGF
DSPTVAYSVLVVAKVEKGKSKKLKSVKELLGITIMERSSFEKNPIDFLEAKGYKEVKKDLIIKLPKYSLFELENGRKRMLASA
GELQKGNELALPSKYVNFLYLASHYEKLKGSPEDNEQKQLFVEQHKHYLDEIIEQISEFSKRVILADANLDKVLSAYNKHR
DKPIREQAENIIHLFTLTNLGAPAAFKYFDTTIDRKRYTSTKEVLDATLIHQSITGLYETRIDLSQLGGD
TLNIEDEYRLHETSKEPDVSLGSTWLSDFPQAWAETGGMGLAVRQAPLIIPLKATSTPVSIKQYPMSQEARLGIKPHIQR
LLDQGILVPCQSPWNTPLLPVKKPGTNDYRPVQDLREVNKRVEDIHPTVPNPYNLLSGLPPSHQWYTVLDLKDAFFCLR
LHPTSQPLFAFEWRDPEMGISGQLTWTRLPQGFKNSPTLFNEALHRDLADFRIQHPDLILLQYVDDLLLAATSELDCQQ
GTRALLQTLGNLGYRASAKKAQICQKQVKYLGYLLKEGQRWLTEARKETVMGQPTPKTPRQLREFLGKAGFCRLFIPGF
AEMAAPLYPLTKPGTLFNWGPDQQKAYQEIKQALLTAPALGLPDLTKPFELFVDEKQGYAKGVLTQKLGPWRRPVAYL
SKKLDPVAAGWPPCLRMVAAIAVLTKDAGKLTMGQPLVILAPHAVEALVKQPPDRWLSNARMTHYQALLLDTDRVQ

FGPVVALNPATLLPLPEEGLQHNCLDILAEAHGTRPDLTDQPLPDADHTWYTDGSSLLQEGQRKAGAAVTTETEVIWA
KALPAGTSAQRAELIALTQALKMAEGKKLNVYTDSRYAFATAHIHGEIYRRRGWLTSEGKEIKNKDEILALLKALFLPKRLS
IIHCPGHQKGHSAEARGNRMADQAARKAAITETPDTSTLLIENSSP

**NFATC2IP-MSH4-PE2 ORF**

AEPVGKRGRWSGGSGAGRGGRGGWGGRGRRPRAQRSPSRGTLDVVSVDLVTDSDEEILEVATARGAADEVEVEPPE
PPGPVASRD GSTGGTGS
QVVSASTCPGTSGAAGDRSSSSSSLPCPAPNSRPAQGSYFGNKRAYAENTVASNFTFGASSSSARDTNYPQTLKTPLST
GNPQRS SSGGSSGGSSGSETPGTSESATPESSGGSSGGSCT
DKKYSIGLDIGTNSVGWAVITDEYKVPSKKFKVLGNTDRHSIKKNLIGALLFDSGETAEATRLKRTARRRYTRRKNRICYLQ
EIFSNEMAKVDDSFFHRLEESFLVEEDKKHERHPIFGNIVDEVAYHEKYPTIYHLRKKLVDSTDKADLRLIYLALAHMIKFR
GHFLIEGDLNPDNSDVDKLFIQLVQTYNQLFEENPINASGVDAKAILSARLSKSRRLENLIAQLPGEKKNGLFGNLIALSLG
LTPNFKSNFDLAEDAKLQLSKDTYDDDLDNLLAQIGDQYADLFLAAKNLSDAILLSDILRVNTEITKAPLSASMIKRYDEHH
QDLTLLKALVRQQLPEKYKEIFFDQSKNGYAGYIDGGASQEEFYKFIKPILEKMDGTEELLVKLNREDLLRKQRTFDNGSIP
HQIHLGELHAILRRQEDFYPFLKDNREKIEKILTFRIPYYVGPLARGNSRFAWMTRKSEETITPWNFEEVVDKGASAQSFI
ERMTNFDKNLPNEKVLPKHSLLYEYFTVYNELTKVKYVTEGMRKPAFLSGEQKKAIVDLLFKTNRKVTVKQLKEDYFKKIE
CFDSVEISGVEDRFNASLGTYHDLLKIIKDKDFLDNEENEDILEDIVLTLTLFEDREMIEERLKTYAHLFDDKVMKQLKRRR
YTGWGRLSRKLINGIRDKQSGKTILDFLKSDGFANRNFMQLIHDDSLTFKEDIQKAQVSGQGDSLHEHIANLAGSPAIKK
GILQTVKVVDELVKVMGRHKPENIVIEMARENQTTQKGQKNSRERMKRIEEGIKELGSQILKEHPVENTQLQNEKLYLY
YLQNGRDMYVDQELDINRLSDYDVDAIVPQSFLKDDSIDNKVLTRSDKNRGKSDNVPSEEVVKKMKNYWRQLLNAKLI
TQRKFDNLTKAERGGLSELDKAGFIKRQLVETRQITKHVAQILDSRMNTKYDENDKLIREVKVITLKSKLVSDFRKDFQFY
KVREINNYHHAHDAYLNAVVGTALIKKYPKLESEFVYGDYKVYDVRKMIAKSEQEIGKATAKYFFYSNIMNFFKTEITLAN
GEIRKRPLIETNGETGEIVWDKGRDFATVRKVLSMPQVNIVKKTEVQTGGFSKESILPKRNSDKLIARKKDWDPKKYGGF
DSPTVAYSVLVVAKVEKGKSKKLKSVKELLGITIMERSSFEKNPIDFLEAKGYKEVKKDLIIKLPKYSLFELENGRKRMLASA
GELQKGNELALPSKYVNFLYLASHYEKLKGSPEDNEQKQLFVEQHKHYLDEIIEQISEFSKRVILADANLDKVLSAYNKHR
DKPIREQAENIIHLFTLTNLGAPAAFKYFDTTIDRKRYTSTKEVLDATLIHQSITGLYETRIDLSQLGGD
TLNIEDEYRLHETSKEPDVSLGSTWLSDFPQAWAETGGMGLAVRQAPLIIPLKATSTPVSIKQYPMSQEARLGIKPHIQR
LLDQGILVPCQSPWNTPLLPVKKPGTNDYRPVQDLREVNKRVEDIHPTVPNPYNLLSGLPPSHQWYTVLDLKDAFFCLR
LHPTSQPLFAFEWRDPEMGISGQLTWTRLPQGFKNSPTLFNEALHRDLADFRIQHPDLILLQYVDDLLLAATSELDCQQ
GTRALLQTLGNLGYRASAKKAQICQKQVKYLGYLLKEGQRWLTEARKETVMGQPTPKTPRQLREFLGKAGFCRLFIPGF
AEMAAPLYPLTKPGTLFNWGPDQQKAYQEIKQALLTAPALGLPDLTKPFELFVDEKQGYAKGVLTQKLGPWRRPVAYL
SKKLDPVAAGWPPCLRMVAAIAVLTKDAGKLTMGQPLVILAPHAVEALVKQPPDRWLSNARMTHYQALLLDTDRVQ
FGPVVALNPATLLPLPEEGLQHNCLDILAEAHGTRPDLTDQPLPDADHTWYTDGSSLLQEGQRKAGAAVTTETEVIWA
KALPAGTSAQRAELIALTQALKMAEGKKLNVYTDSRYAFATAHIHGEIYRRRGWLTSEGKEIKNKDEILALLKALFLPKRLS
IIHCPGHQKGHSAEARGNRMADQAARKAAITETPDTSTLLIENSSP

**NFATC2IP-CDKN2A-PE2 ORF**

AEPVGKRGRWSGGSGAGRGGRGGWGGRGRRPRAQRSPSRGTLDVVSVDLVTDSDEEILEVATARGAADEVEVEPPE
PPGPVASRD GSTGGTGS
EPAAGSSMEPSADWLATAAARGRVEEVRALLEAGALPNAPNSYGRRPIQVMMMGSARVAELLLLHGAEPNCADPAT
LTRPVHDAA SSGGSSGGSSGSETPGTSESATPESSGGSSGGSCT
DKKYSIGLDIGTNSVGWAVITDEYKVPSKKFKVLGNTDRHSIKKNLIGALLFDSGETAEATRLKRTARRRYTRRKNRICYLQ
EIFSNEMAKVDDSFFHRLEESFLVEEDKKHERHPIFGNIVDEVAYHEKYPTIYHLRKKLVDSTDKADLRLIYLALAHMIKFR
GHFLIEGDLNPDNSDVDKLFIQLVQTYNQLFEENPINASGVDAKAILSARLSKSRRLENLIAQLPGEKKNGLFGNLIALSLG

LTPNFKSNFDLAEDAKLQLSKDTYDDDLDNLLAQIGDQYADLFLAAKNLSDAILLSDILRVNTEITKAPLSASMIKRYDEHH
QDLTLLKALVRQQLPEKYKEIFFDQSKNGYAGYIDGGASQEEFYKFIKPILEKMDGTEELLVKLNREDLLRKQRTFDNGSIP
HQIHLGELHAILRRQEDFYPFLKDNREKIEKILTFRIPYYVGPLARGNSRFAWMTRKSEETITPWNFEEVVDKGASAQSFI
ERMTNFDKNLPNEKVLPKHSLLYEYFTVYNELTKVKYVTEGMRKPAFLSGEQKKAIVDLLFKTNRKVTVKQLKEDYFKKIE
CFDSVEISGVEDRFNASLGTYHDLLKIIKDKDFLDNEENEDILEDIVLTLTLFEDREMIEERLKTYAHLFDDKVMKQLKRRR
YTGWGRLSRKLINGIRDKQSGKTILDFLKSDGFANRNFMQLIHDDSLTFKEDIQKAQVSGQGDSLHEHIANLAGSPAIKK
GILQTVKVVDELVKVMGRHKPENIVIEMARENQTTQKGQKNSRERMKRIEEGIKELGSQILKEHPVENTQLQNEKLYLY
YLQNGRDMYVDQELDINRLSDYDVDAIVPQSFLKDDSIDNKVLTRSDKNRGKSDNVPSEEVVKKMKNYWRQLLNAKLI
TQRKFDNLTKAERGGLSELDKAGFIKRQLVETRQITKHVAQILDSRMNTKYDENDKLIREVKVITLKSKLVSDFRKDFQFY
KVREINNYHHAHDAYLNAVVGTALIKKYPKLESEFVYGDYKVYDVRKMIAKSEQEIGKATAKYFFYSNIMNFFKTEITLAN
GEIRKRPLIETNGETGEIVWDKGRDFATVRKVLSMPQVNIVKKTEVQTGGFSKESILPKRNSDKLIARKKDWDPKKYGGF
DSPTVAYSVLVVAKVEKGKSKKLKSVKELLGITIMERSSFEKNPIDFLEAKGYKEVKKDLIIKLPKYSLFELENGRKRMLASA
GELQKGNELALPSKYVNFLYLASHYEKLKGSPEDNEQKQLFVEQHKHYLDEIIEQISEFSKRVILADANLDKVLSAYNKHR
DKPIREQAENIIHLFTLTNLGAPAAFKYFDTTIDRKRYTSTKEVLDATLIHQSITGLYETRIDLSQLGGD
TLNIEDEYRLHETSKEPDVSLGSTWLSDFPQAWAETGGMGLAVRQAPLIIPLKATSTPVSIKQYPMSQEARLGIKPHIQR
LLDQGILVPCQSPWNTPLLPVKKPGTNDYRPVQDLREVNKRVEDIHPTVPNPYNLLSGLPPSHQWYTVLDLKDAFFCLR
LHPTSQPLFAFEWRDPEMGISGQLTWTRLPQGFKNSPTLFNEALHRDLADFRIQHPDLILLQYVDDLLLAATSELDCQQ
GTRALLQTLGNLGYRASAKKAQICQKQVKYLGYLLKEGQRWLTEARKETVMGQPTPKTPRQLREFLGKAGFCRLFIPGF
AEMAAPLYPLTKPGTLFNWGPDQQKAYQEIKQALLTAPALGLPDLTKPFELFVDEKQGYAKGVLTQKLGPWRRPVAYL
SKKLDPVAAGWPPCLRMVAAIAVLTKDAGKLTMGQPLVILAPHAVEALVKQPPDRWLSNARMTHYQALLLDTDRVQ
FGPVVALNPATLLPLPEEGLQHNCLDILAEAHGTRPDLTDQPLPDADHTWYTDGSSLLQEGQRKAGAAVTTETEVIWA
KALPAGTSAQRAELIALTQALKMAEGKKLNVYTDSRYAFATAHIHGEIYRRRGWLTSEGKEIKNKDEILALLKALFLPKRLS
IIHCPGHQKGHSAEARGNRMADQAARKAAITETPDTSTLLIENSSP

**NFATC2IP-PE2 ORF**

AEPVGKRGRWSGGSGAGRGGRGGWGGRGRRPRAQRSPSRGTLDVVSVDLVTDSDEEILEVATARGAADEVEVEPPE
PPGPVASRD SSGGSSGGSSGSETPGTSESATPESSGGSSGGSCT
DKKYSIGLDIGTNSVGWAVITDEYKVPSKKFKVLGNTDRHSIKKNLIGALLFDSGETAEATRLKRTARRRYTRRKNRICYLQ
EIFSNEMAKVDDSFFHRLEESFLVEEDKKHERHPIFGNIVDEVAYHEKYPTIYHLRKKLVDSTDKADLRLIYLALAHMIKFR
GHFLIEGDLNPDNSDVDKLFIQLVQTYNQLFEENPINASGVDAKAILSARLSKSRRLENLIAQLPGEKKNGLFGNLIALSLG
LTPNFKSNFDLAEDAKLQLSKDTYDDDLDNLLAQIGDQYADLFLAAKNLSDAILLSDILRVNTEITKAPLSASMIKRYDEHH
QDLTLLKALVRQQLPEKYKEIFFDQSKNGYAGYIDGGASQEEFYKFIKPILEKMDGTEELLVKLNREDLLRKQRTFDNGSIP
HQIHLGELHAILRRQEDFYPFLKDNREKIEKILTFRIPYYVGPLARGNSRFAWMTRKSEETITPWNFEEVVDKGASAQSFI
ERMTNFDKNLPNEKVLPKHSLLYEYFTVYNELTKVKYVTEGMRKPAFLSGEQKKAIVDLLFKTNRKVTVKQLKEDYFKKIE
CFDSVEISGVEDRFNASLGTYHDLLKIIKDKDFLDNEENEDILEDIVLTLTLFEDREMIEERLKTYAHLFDDKVMKQLKRRR
YTGWGRLSRKLINGIRDKQSGKTILDFLKSDGFANRNFMQLIHDDSLTFKEDIQKAQVSGQGDSLHEHIANLAGSPAIKK
GILQTVKVVDELVKVMGRHKPENIVIEMARENQTTQKGQKNSRERMKRIEEGIKELGSQILKEHPVENTQLQNEKLYLY
YLQNGRDMYVDQELDINRLSDYDVDAIVPQSFLKDDSIDNKVLTRSDKNRGKSDNVPSEEVVKKMKNYWRQLLNAKLI
TQRKFDNLTKAERGGLSELDKAGFIKRQLVETRQITKHVAQILDSRMNTKYDENDKLIREVKVITLKSKLVSDFRKDFQFY
KVREINNYHHAHDAYLNAVVGTALIKKYPKLESEFVYGDYKVYDVRKMIAKSEQEIGKATAKYFFYSNIMNFFKTEITLAN
GEIRKRPLIETNGETGEIVWDKGRDFATVRKVLSMPQVNIVKKTEVQTGGFSKESILPKRNSDKLIARKKDWDPKKYGGF
DSPTVAYSVLVVAKVEKGKSKKLKSVKELLGITIMERSSFEKNPIDFLEAKGYKEVKKDLIIKLPKYSLFELENGRKRMLASA
GELQKGNELALPSKYVNFLYLASHYEKLKGSPEDNEQKQLFVEQHKHYLDEIIEQISEFSKRVILADANLDKVLSAYNKHR
DKPIREQAENIIHLFTLTNLGAPAAFKYFDTTIDRKRYTSTKEVLDATLIHQSITGLYETRIDLSQLGGD

TLNIEDEYRLHETSKEPDVSLGSTWLSDFPQAWAETGGMGLAVRQAPLIIPLKATSTPVSIKQYPMSQEARLGIKPHIQR
LLDQGILVPCQSPWNTPLLPVKKPGTNDYRPVQDLREVNKRVEDIHPTVPNPYNLLSGLPPSHQWYTVLDLKDAFFCLR
LHPTSQPLFAFEWRDPEMGISGQLTWTRLPQGFKNSPTLFNEALHRDLADFRIQHPDLILLQYVDDLLLAATSELDCQQ
GTRALLQTLGNLGYRASAKKAQICQKQVKYLGYLLKEGQRWLTEARKETVMGQPTPKTPRQLREFLGKAGFCRLFIPGF
AEMAAPLYPLTKPGTLFNWGPDQQKAYQEIKQALLTAPALGLPDLTKPFELFVDEKQGYAKGVLTQKLGPWRRPVAYL
SKKLDPVAAGWPPCLRMVAAIAVLTKDAGKLTMGQPLVILAPHAVEALVKQPPDRWLSNARMTHYQALLLDTDRVQ
FGPVVALNPATLLPLPEEGLQHNCLDILAEAHGTRPDLTDQPLPDADHTWYTDGSSLLQEGQRKAGAAVTTETEVIWA
KALPAGTSAQRAELIALTQALKMAEGKKLNVYTDSRYAFATAHIHGEIYRRRGWLTSEGKEIKNKDEILALLKALFLPKRLS
IIHCPGHQKGHSAEARGNRMADQAARKAAITETPDTSTLLIENSSP

**IGF-PE2 ORF**

REGMNKPEGYGEEERRAPQEGIVDECCFRECDLRRLEMYCAPLKPAKEAREVRAQRHEDMPKEQKYQPPEENKNEKE
QRRKGEEF <u>SSGGSSGGSSGSETPGTSESATPESSGGSSGGSCT</u>
DKKYSIGLDIGTNSVGWAVITDEYKVPSKKFKVLGNTDRHSIKKNLIGALLFDSGETAEATRLKRTARRRYTRRKNRICYLQ
EIFSNEMAKVDDSFFHRLEESFLVEEDKKHERHPIFGNIVDEVAYHEKYPTIYHLRKKLVDSTDKADLRLIYLALAHMIKFR
GHFLIEGDLNPDNSDVDKLFIQLVQTYNQLFEENPINASGVDAKAILSARLSKSRRLENLIAQLPGEKKNGLFGNLIALSLG
LTPNFKSNFDLAEDAKLQLSKDTYDDDLDNLLAQIGDQYADLFLAAKNLSDAILLSDILRVNTEITKAPLSASMIKRYDEHH
QDLTLLKALVRQQLPEKYKEIFFDQSKNGYAGYIDGGASQEEFYKFIKPILEKMDGTEELLVKLNREDLLRKQRTFDNGSIP
HQIHLGELHAILRRQEDFYPFLKDNREKIEKILTFRIPYYVGPLARGNSRFAWMTRKSEETITPWNFEEVVDKGASAQSFI
ERMTNFDKNLPNEKVLPKHSLLYEYFTVYNELTKVKYVTEGMRKPAFLSGEQKKAIVDLLFKTNRKVTVKQLKEDYFKKIE
CFDSVEISGVEDRFNASLGTYHDLLKIIKDKDFLDNEENEDILEDIVLTLTLFEDREMIEERLKTYAHLFDDKVMKQLKRRR
YTGWGRLSRKLINGIRDKQSGKTILDFLKSDGFANRNFMQLIHDDSLTFKEDIQKAQVSGQGDSLHEHIANLAGSPAIKK
GILQTVKVVDELVKVMGRHKPENIVIEMARENQTTQKGQKNSRERMKRIEEGIKELGSQILKEHPVENTQLQNEKLYLY
YLQNGRDMYVDQELDINRLSDYDVDAIVPQSFLKDDSIDNKVLTRSDKNRGKSDNVPSEEVVKKMKNYWRQLLNAKLI
TQRKFDNLTKAERGGLSELDKAGFIKRQLVETRQITKHVAQILDSRMNTKYDENDKLIREVKVITLKSKLVSDFRKDFQFY
KVREINNYHHAHDAYLNAVVGTALIKKYPKLESEFVYGDYKVYDVRKMIAKSEQEIGKATAKYFFYSNIMNFFKTEITLAN
GEIRKRPLIETNGETGEIVWDKGRDFATVRKVLSMPQVNIVKKTEVQTGGFSKESILPKRNSDKLIARKKDWDPKKYGGF
DSPTVAYSVLVVAKVEKGKSKKLKSVKELLGITIMERSSFEKNPIDFLEAKGYKEVKKDLIIKLPKYSLFELENGRKRMLASA
GELQKGNELALPSKYVNFLYLASHYEKLKGSPEDNEQKQLFVEQHKHYLDEIIEQISEFSKRVILADANLDKVLSAYNKHR
DKPIREQAENIIHLFTLTNLGAPAAFKYFDTTIDRKRYTSTKEVLDATLIHQSITGLYETRIDLSQLGGD
TLNIEDEYRLHETSKEPDVSLGSTWLSDFPQAWAETGGMGLAVRQAPLIIPLKATSTPVSIKQYPMSQEARLGIKPHIQR
LLDQGILVPCQSPWNTPLLPVKKPGTNDYRPVQDLREVNKRVEDIHPTVPNPYNLLSGLPPSHQWYTVLDLKDAFFCLR
LHPTSQPLFAFEWRDPEMGISGQLTWTRLPQGFKNSPTLFNEALHRDLADFRIQHPDLILLQYVDDLLLAATSELDCQQ
GTRALLQTLGNLGYRASAKKAQICQKQVKYLGYLLKEGQRWLTEARKETVMGQPTPKTPRQLREFLGKAGFCRLFIPGF
AEMAAPLYPLTKPGTLFNWGPDQQKAYQEIKQALLTAPALGLPDLTKPFELFVDEKQGYAKGVLTQKLGPWRRPVAYL
SKKLDPVAAGWPPCLRMVAAIAVLTKDAGKLTMGQPLVILAPHAVEALVKQPPDRWLSNARMTHYQALLLDTDRVQ
FGPVVALNPATLLPLPEEGLQHNCLDILAEAHGTRPDLTDQPLPDADHTWYTDGSSLLQEGQRKAGAAVTTETEVIWA
KALPAGTSAQRAELIALTQALKMAEGKKLNVYTDSRYAFATAHIHGEIYRRRGWLTSEGKEIKNKDEILALLKALFLPKRLS
IIHCPGHQKGHSAEARGNRMADQAARKAAITETPDTSTLLIENSSP
